# collectNET: a web server for integrated inference of cell-cell communication network

**DOI:** 10.1101/2024.03.18.585493

**Authors:** Yan Pan, Zijing Gao, Xuejian Cui, Zhen Li, Rui Jiang

**Affiliations:** Ministry of Education Key Laboratory of Bioinformatics, Bioinformatics Division at the Beijing National Research Center for Information Science and Technology, Center for Synthetic and Systems Biology, Department of Automation, Tsinghua University, Beijing 100084, China

## Abstract

**Summary:** Cell-cell communication through ligand-receptor pairs forms the cornerstone for complex functionalities in multicellular organisms. Deciphering such intercellular signaling can contribute to unraveling disease mechanisms and enables targeted therapy. Nonetheless, notable biases and inconsistencies are evident among the inferential outcomes generated by current methods for inferring cell-cell communication network. To fill this gap, we developed collectNET (http://health.tsing-hua.edu.cn/collectnet) as the first web server for efficiently inferring the cell-cell communication network, with efficient calculation, hierarchical browsing, comprehensive statistics, advanced searching, and intuitive visualization. collectNET provides a reliable online inference service with prior knowledge of three public ligand-receptor databases and systematic integration of three mainstream inference methods. Additionally, collectNET has assembled a human cell-cell communication atlas, including 126,785 significant communication pairs based on 343,023 single cells. We anticipate that collectNET will benefit researchers in gaining a more holistic understanding of cell development and differentiation mechanisms.

**Availability and Implementation:** collectNET is freely available at http://health.tsinghua.edu.cn/collectnet.

**Supplementary information:** Supplementary data are available at *Bioinformatics* online.

## 1 Introduction

Cell-cell communication (CCC) forms the cornerstone of life activities in multicellular organisms, involving the interaction between ligand-receptor (L-R) pairs (Almet, et al., 2021; Song, et al., 2019). Ligands are proteins that bind to other biomolecules to transmit signals, while receptors are biological molecules that interact with ligands (Armingol, et al., 2021). Binding of ligands to receptors on recipient cell membranes triggers the activation of transcription factors and their target genes within the cell (Ramilowski, et al., 2015; Wang, Almet and Nie, 2023). In this manner, cell-cell communication orchestrates processes such as cell proliferation, migration, differentiation, and death, thereby achieving tissue-wide coordination, ultimately resulting in the emergence of complex functionality in cellular organisms. Therefore, deciphering the cell-cell communication network plays a pivotal role in advancing the understanding of cell development, tissue homeostasis, immune processes, and further applied to the diagnosis and treatment of major diseases such as cancer (Peng, et al., 2022).

Recent advances in single-cell RNA sequencing (scRNA-seq) technology and the accumulation of scRNA-seq data enable the measurement of ligand and receptor expression in single-cells across various cell types, thereby assisting in the inference and construction of cell-cell communication networks, further enhancing the understanding of human disease mechanisms.

Previous methods employ various approaches to calculate the communication scores between ligand and receptor (Cillo, et al., 2020; Efremova, et al., 2020; Jin, et al., 2021). These methods follow a common computational framework, where feature genes associated with cell-cell communication are selected from the matrix. Subsequently, communication scores (i.e., interaction strengths) of ligand-receptor pairs are computed. These scores are further utilized to estimate the communication status between cells along each pathway (Peng, et al., 2022; Wang, Almet and Nie, 2023). For example, CellTalker (Cillo, et al., 2020) uses ligands and receptors that exhibit non-zero expression in a predefined proportion of cells. Cell-Chat (Jin, et al., 2021) calculates communication scores by multiplying the gene expression of ligands and receptors, which take effects by taking the geometric average of polymer-containing ligands or receptors before the multiplication. CellPhoneDB (Efremova, et al., 2020) has proposed enrichment scores, which represent the minimum average gene expression of receptors in specific cell types. Nonetheless, previous benchmarking studies have observed significant differences between the results of these methods (Dimitrov, et al., 2022; Liu, Sun and Wang, 2022). For example, LIANA found that regardless of variations in inference methods or reference databases, the degree of overlap remained relatively low within the same dataset (Dimitrov, et al., 2022). In addition, there remains a deficiency in comprehensive knowledge representation of the cell-cell communication patterns in human organs. While some databases or web servers partially address cell-cell communication and provide lists of inferred results (He, et al., 2020; Ma, et al., 2024; Pan, et al., 2023), many of these web-servers or databases lack online computational services, requiring users to have local computing resources and deploy development environments. Since existing methods vary in their computational resource requirements, users may also find it difficult to complete the inference and analysis on ordinary computers. Additionally, there are shortcomings in the visualization and analysis of inferred cell communication network results (Supplementary Table S1). Therefore, an integrated network platform for inferring, analyzing, and visualizing cell communication networks is in pressing need.

To circumvent these bottlenecks, we develop collectNET (Combined ceLL-cEll CommunicaTion NETwork), a comprehensive web-platform for efficiently inference of cell-cell communication network and better interrogating cell-cell communication networks across multiple human organs (Figure 1). With prior knowledge of 3,954 ligand-receptor pairs from three public databases and systematic integration of three widely-used inference methods, collectNET provides a comprehensive and reliable online inference workflow, including data pre-processing, inference, and visualization, and an atlas of cell-cell communication across human organs with 126,785 significant communication pairs on 31 organs and 485 cell types based on the scRNA-seq datasets of 343,023 single cells. With ample computational services, collectNET eliminates the need for any local computational storage or resources. The inferential outcomes of cell-cell communication can be used to identify the topological characteristics and communication pattern features, thereby providing a theoretical foundation for exploring disease mechanisms and conducting precision medicine research. We anticipate that collectNET will demonstrate its extensive applicability in elucidating cell development and differentiation mechanisms.

**Fig. 1.**
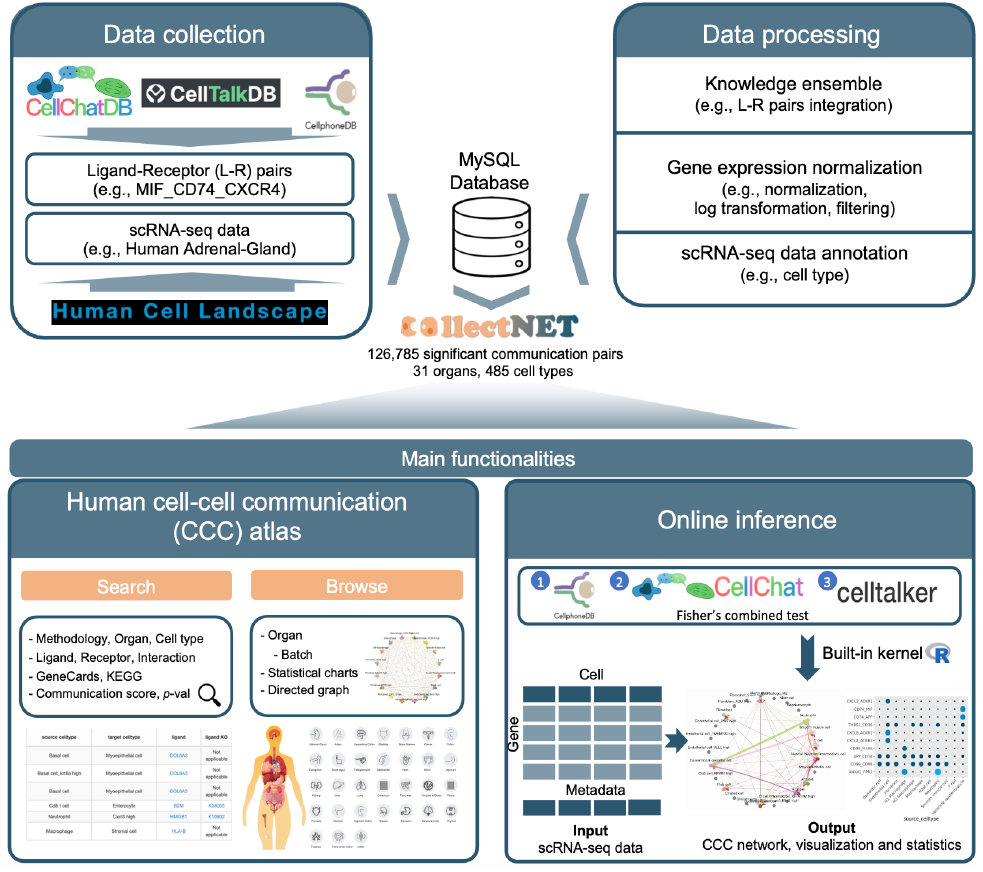
Overview of the collectNET website. collectNET is a web server for integrated inference of cell-cell communication network, with efficient calculation, hierarchical browsing, comprehensive statistics, advanced search capabilities, and intuitive visualization.

## 2 Materials, methods and web interface

We first obtained a more comprehensive ligand-receptor pair database by integrating multiple databases, namely CellPhoneDB (Efremova, et al., 2020), CellTalkDB (Shao, et al., 2021), and CellChatDB (Jin, et al., 2021). The database comprises 3,954 ligand-receptor pairs, including complexes and polymers (Supplementary Table S2). In the construction of the cell-cell communication network atlas, we utilized single-cell data of 31 human organs from the Human Cell Landscape (HCL) (Han, et al., 2020) as the raw count gene expression matrix input and conducted a series of preprocessing steps upon them (Supplementary Text S1). Then, these datasets underwent preprocessing steps, including normalization, logarithmic transformation, and filtering out genes expressed in fewer than 3 cells and cells expressing fewer than 200 genes.

Then, we utilized an integration approach to complete the online inference (Supplementary Text S2). Firstly, we referred to the modeling and communication pattern analysis methods for cell-cell communication in the first and second studies and utilized Fisher’s combined probability test (Fisher, 1955) to integrate the results of multiple cell communication networks and combined both the *p*-value matrix and the normalized gene count matrix. Specifically, if all the null hypotheses are true and the p-values are independent of each other, the sum of the logarithm of their test statistics follows a chi-square distribution with 2*k* degrees of freedom, where *k* is the number of tests to be merged.

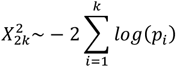

Consequently, by integrating three mutually independent hypothesis tests conducted for the interactions between ligands and receptors, we obtained the statistical significance of considering multiple individual experiments as a whole. Additionally, for the raw count gene expression matrix, we first normalized each matrix individually by linearly mapping the values to a range between 0 and 1, then summed and normalized the corresponding indexed values in the same way to obtain the averaged communication score matrix.

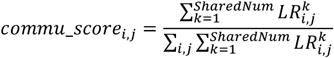

Here, *i* and *j* denote the index of corresponding cell types, and *S*h*aredNum* denotes how many ligand-receptor pairs appear in the pair library. After normalization, we obtained communication scores for each ligand-receptor pair across various cell types and subsequently filtered out ligand-receptor pairs that have values lower than a specific threshold in the corresponding positions of the *p*-value matrix.

Taken together, collectNET was developed using MySQL, PHP, JavaScript, and HTML (Supplementary Text S5), and it comprises seven main pages offering extensive interfaces (Supplementary Text S3). On the *Search* page, users can select significant receptor-ligand pair s through advanced options. On the *Browse* page, users can hierarchically query L-R pairs for specific organs and browse various statistical measures. On the *Analysis* page, users can effortlessly perform cell communication network inference without programming. On the *Download* page, users can download all the necessary information in the atlas. On the pages such as *Home, Help*, and *About*, users can find tutorials and other information about the usage of the website.

## 3 Functionalities and Applications

The major functionalities of collectNET encompass interrogation, online inference, visualization and analysis of cell-cell communication (Supplementary Text S3).

### Intuitively browsing

collectNET is meticulously organized according to organ systems, offering users a comprehensive and visually engaging experience. For each organ, collectNET presents a nuanced depiction of cellular composition through circular charts that enumerate the cell counts for different cell types. Additionally, collectNET provides a detailed account of the most frequently occurring ligands, receptors, and ligand-receptor pairs, as identified by our cell-cell communication inference methodology, along with their respective frequencies, in a tabular format that underscores the significance of these interactions.

### Advanced searching

collectNET offers advanced search capabilities that facilitate comprehensive exploration of biological interactions. Users have the flexibility to index and query the database using various parameters such as methods, organs, cell types, receptors, ligands, or ligand-receptor interactions, with multi-criteria indexing supported.

### Online inference and analysis

For online analysis, two input files are required: one is a file containing the single-cell gene expression matrix, and the other is a file containing the cell annotations. We also offer users options to specify various parameters (Supplementary Table S3). Under typical dataset scales, the computation time ranges from several minutes to a dozen minutes (Supplementary Fig. S4). The inference results include detailed inference results and figures of the most frequently occurring ligands and receptors.

In addition to the functionalities mentioned above, collectNET has demonstrated a diverse range of application scenarios (Supplementary Text S4; Supplementary Figs. S1-S3). Here, we describe three main application scenarios.

In the first scenario, a researcher aims to investigate the cell-cell communication signaling pathways in which a specific gene is involved, as well as the likelihood of its involvement in cell-cell communication across human organs. collectNET can substantially aid in this endeavor (Supplementary Fig. S1). In specific, the researcher can firstly select the name of the ligand or receptor and select to sort the search results by inference *p*-value or communication score in descending order on the *Search* page. Then, from the search outcomes, the researcher can discern the significant cell-cell communications associated with the gene, occurring in various cell types, ranked by communication strength or confidence level. Furthermore, the researcher can access additional information such as annotations for each ligand-receptor pair in external databases, along with other pertinent details, and download these data. For example, if the researcher has a particular interest in the communication status within human heart, the *Browse* page of collectNET articulates a comprehensive array of cell types encapsulated within that organ. Predominantly, CD74 emerges as the most frequently occurring ligand, with an incidence of 142 occurrences, while MIF stands out as the recurrent receptor, noted 89 times. The most prevalent ligand-receptor pair is identified as CD74_MIF. Within the cardiac dataset, delineated into two distinct batches, one can procure the inferential outcomes garnered from three disparate methodologies, alongside the visualizations of inferences culled from collectNET, which articulate the count of significant ligand-receptor pairs among various cell types in conjunction with the composite communication scores. An explication of the more intricate details of the graph is furnished in the tabular form beneath the visualizations.

In the second scenario, if users aim to explore the internal structure of communication networks, this can be achieved by exploring the topological characteristics of the network. In the tutorial data provided in collectNET (Supplementary Fig. S3), the ventricle cardiomyocyte exhibits a PageRank score approaching 0.2, denoting a pronounced importance in the tutorial’s cardiac data. This corresponds to the high attention paid to the ventricle cardiomyocytes as a specialized class of myocytes that are uniquely found within the ventricles of the heart (Ahuja, Sdek and MacLellan, 2007). In contrast, dendritic cells, endothelial cells, and neutrophils all exhibit significantly lower scores, suggesting a more singular signaling pathway presence within these cell types. Also, ventricle cardiomyocytes, macrophages, fibroblasts, smooth muscle cells, M1 macrophages, and M2 macrophages can be identified as the largest clique within the communication network. This observation suggests that these cell types may share a substantial number of common communication patterns, exhibit highly active interactions among each other, and should be comprehensively analyzed within the entire clique for communication studies. Additionally, by performing graph clustering on the communication network given by collectNET, it is evident that, in addition to the aforementioned largest clique, two distinct clusters have emerged: one comprising endothelial cells and neutrophils, and the other dendritic cells. The discovery can be justified since both endothelial cells and neutrophils are associated with the vascular and blood systems, interacting closely in inflammation and immune responses, such as the release of inflammatory mediators and neutrophil migration (Phillipson and Kubes, 2011; Tonnesen, 1989). This clustering approach provides insights into the functional relationships and potential collaborative roles of these cell types in the context of cardiovascular and immunological processes.

Furthermore, we have validated the online computation superiority and robustness. We calculated Pearson correlation coefficients (Cohen, et al., 2009) on the communication networks inferred by the four methods across ten independent single-cell datasets from HCL (Supplementary Text S4 and Supplementary Fig. S2). Through a boxplot of the Pearson correlation coefficient between every single method and the other methods, it is evident that the results of collectNET are highly correlated to those of each method, with the median value approaching 0.8, 0.9 and 1.0 respectively. This underscores that despite the obvious differences between the networks inferred by different methods, collectNET effectively integrates the methods based on different models and databases of L-R pairs, constructing a more accurate communication network by combining diverse biological information.

## 4 Conclusion and Discussion

Cell-cell communication is accomplished through L-R pairs based inter-cellular signaling. The study of cell-cell communication is of great significance as it allows for the investigation of fundamental biological questions such as development and differentiation, the understanding of how overall tissue coordination and functionality are achieved, as well as the analysis of the coordinated actions of multiple cells in disease onset. Currently, various methods have been developed to infer cell-cell communication from single-cell transcriptomic sequencing data, but significant biases and inconsistencies in the inference results of these methods remain to be shown. collectNET is an integrated online tool for cell-cell communication analysis along with a human cell-cell communication atlas.

Compared with existing web-tools for cell-cell communication, collectNET highlights five notable features: (i) collectNET integrates CellChat (Jin, et al., 2021), CellPhoneDB (Efremova, et al., 2020) and CellTalker (Cillo, et al., 2020) based on Fisher’s combined probability test, thereby enhancing the reliability of intercellular communication inference; (ii) collectNET incorporates three widely-used databases of ligand-receptor pairs as prior knowledge to extract maximum information from the input data; (iii) collectNET enables users lacking coding expertise or computational resources to perform efficient inference and visualization through intuitive tutorial-style interfaces; (iv) collectNET provides downloadable tables for diverse methods, organs, and user-submitted tasks, supporting versatile applications like drug target prediction and immunotherapy strategies; (v) collectNET furnishes extensive cell-cell communication resource, encompassing 126,785 significant communication pairs on 31 organs and 485 cell types based on the scRNA-seq datasets of 343,023 single cells.

In future versions, collectNET will undergo further improvements and enhancements in the following two aspects. Firstly, we will keep pace with updates of various cell-cell communication inference methods to ensure the completeness and accuracy of the inference results. Furthermore, the integration of a diverse array of omics data, including single-cell spatial omics data, will be contemplated. This approach will introduce coordinate information into the inference of cell-cell communications, thereby enhancing the precision and reliability of the derived interactions. Such a multidimensional expansion of data sources is expected to provide a more nuanced and contextually grounded representation of cellular dialogues within the biological system. We anticipate that collectNET will benefit both biologists and algorithm researchers in better understanding cellular differentiation, developmental mechanisms, and exploring strategies for disease treatment.

