## Supplementary information for "collectNET: a web server for integrated inference of cell-cell communication network"

### Supplementary Texts

#### Text S1. Data collection and processing

##### Data collection and processing

To fully harness the potential ligand-receptor information from single-cell datasets, we first obtained a more comprehensive and complete ligand-receptor pair database by integrating multiple databases. In collectNET, three widely-used public ligand-receptor pair databases, namely CellPhoneDB (Efremova, et al., 2020), CellTalkDB (Shao, et al., 2021), and CellChatDB (Jin, et al., 2021), are integrated to form a database comprising 3,954 ligand-receptor pairs, including complexes and polymers.

Human Cell Landscape (HCL) is a comprehensive and publicly available human single-cell data atlas (Han, et al., 2020). In the construction of the cell-cell communication network atlas, we utilized single-cell data from the HCL database as the input. First, we collected 31 main human organ datasets from the HCL database, comprising a total of 485 cell types and 343,023 cells. Then, these datasets underwent preprocessing steps. Initially, they were normalized using the method described below. Here, $X_{i,j}$ denotes the expression level of the $i$-th gene in the $j$-th cell from the input raw count matrix, $lb_j$ represents the library size of the j-th cell, and $sf_j$ signifies the size factor for the j-th cell. $X_{norm\ i,j}$ is the normalized expression level of the $i$-th gene in the $j$-th cell.

$$lb_j = \sum_{i=1}^{N} X_{i,j}\,, \qquad sf_j = \frac{lb_j}{\overline{lb}}, \qquad X_{norm\ i,j} = \frac{X_{i,j}}{sf_j}$$

Then logarithmic transformation is performed on the normalized gene expression matrix. Here $X_{log\ i,j}$ is the log-transformed expression level of the $i$-th gene in the $j$-th cell.

$$X_{log\ i,j} = log_2\left(1 + X_{norm\ i,j}\right)$$

During the construction process of the cell-cell communication atlas, genes expressed in fewer than 3 cells and cells expressing fewer than 200 genes were filtered out. In the online inference interface, users also have the option to customize the parameters according to their preferences.

#### Statistics and visualization

To enhance user-friendliness, collectNET presents information within the atlas through a variety of statistical metrics and visualizations. We have quantified the number of cells for each cell type within every human organ, and identified the top 10 ligands, receptors, and ligand-receptor pairs that appear most frequently across all significant pathways, which are then visualized using circular slice charts and histograms. For each dataset within individual human organs, we display the results from CellTalker (Cillo, et al., 2020), CellPhoneDB (Efremova, et al., 2020), and CellChat (Jin, et al., 2021) during the integrated inference process, as well as the number and average communication score of significant ligand-receptor pairs between each pair of cell types after integration. In the table titled "Significant Ligand-Receptor Pairs" at the bottom, we provide detailed information including the inference method, organ, batch, source and target cell types, ligand, receptor, ligand-receptor pair, communication score, and $p$-value of the Fisher combined probability test in the inference method.

In order to elucidate the biological significance of various receptors in signaling pathways, we provide GeneCards (Safran, et al., 2010) and Kyoto Encyclopedia of Genes and Genomes (KEGG) (Kanehisa and Goto, 2000) links for ligands and receptors in the tables as well. The GeneCards links primarily emphasize the characteristics of the genes themselves, including their HUGO Gene Nomenclature Committee (HGNC) symbols, GeneCards identifiers, enhancers, and silencers. The KEGG links, on the other hand, direct to the orthology of the corresponding ligands or receptors, providing information on potentially related genes, signaling pathways, and diseases. These reference links provide insights into the biological background of the receptors and their potential pathological associations, enriching the knowledge graph of the communication network.

#### Text S2. Methodology for online inference

After obtaining the integrated ligand-receptor database and single-cell transcriptome dataset, we employed these two inputs for online inference of cell-cell communication. Due to the different statistical models underlying distinct inference methods, the robustness against noise varies. Therefore, we utilized an integration approach to extract relevant information from the inference results.

Firstly, we referred to the modeling and communication pattern analysis methods for cell-cell communication in the first and second studies and utilized Fisher's combined probability test to integrate the results of multiple cell communication networks (Fisher, 1955), thereby achieving more precise construction of the communication graph. In the integration approach, our goal was to combine both the $p$-value matrix and the communication strength matrix for each single method. Here the $p$-value matrix represents the significance of whether a certain ligand-receptor pair expresses between two cell types, and the communication strength matrix represents the relative interacting strength for a certain ligand-receptor pair. For the integration of the $p$-value matrix, Fisher's combined probability test (Fisher, 1955) was employed since multiple independent tests are based on the same null hypothesis of an interacting pair not existing between two cell types. Specifically, if all the null hypotheses are true and the $p$-value of each single test is independent of each other, the sum of the logarithm of their test statistics follows a chi-square distribution with $2k$ degrees of freedom, where $k$ is the number of tests to be merged. The formula is as follows:

$$X_{2k}^{2} \sim -2\sum_{i=1}^{k} log(p_i)$$

In this equation, since we employed three methods for integration, k is set to 3. Consequently, by integrating three mutually independent hypothesis tests conducted on the interactions between ligands and receptors, we obtained $p$-values with an overall significance. Additionally, for the three normalized communication strength matrix, we first normalize each matrix individually by linearly mapping the values to a range between 0 and 1, then summed and normalized the corresponding indexed values in the same way to obtain the averaged communication score matrix between cell types.

$$commu\_score_{i,j} = \frac{\sum_{k=1}^{SharedNum} LR_{i,j}^{k}}{\sum_{i,j}\sum_{k=1}^{SharedNum} LR_{i,j}^{k}}$$

Here, $i$ and $j$ denote the corresponding cell types, $SharedNum$ denotes the number of ligand-receptor pairs appear in the pair library. The average operation was applied in this context because it is not affected by zero expression levels of certain ligand-receptor pairs or the gene expression nature of sparsity. After normalization, we obtained communication scores for each ligand-receptor pair across various cell types. Here, communication score is a normalized value ranging from 0 to 1 that represents the strength of communication between a pair of ligand-receptor. We then filtered out ligand-receptor pairs that have values lower than a specific threshold in the corresponding positions of the $p$-value matrix. The threshold value is set at 0.05, and we regard these ligand-receptor pairs as significant in the respective cell type pairs.

#### Text S3. Web interface for collectNET

collectNET includes seven main pages to provide convenient and efficient online analysis along with clear visualization of the results. The *Home* page highlights the key features and major applications of collectNET. The *Browse* page presents hierarchical cell-cell communication networks and tables for multiple batches of human organs, which can be searched item by item on the *Search* page. The *Download* page provides access to all the data encompassed in the human cell-cell communication atlas. Detailed instructions regarding the usage of each page and frequently asked questions are provided in the *Help* page, and high-lighted information of collectNET can be found on the *About* page.

**Input**

On the *Analysis* page, the online cell communication network inference functionality of collectNET can be utilized. Two input files are required: one is a file containing the single-cell gene expression matrix, and the other is a comma-separated values file (csv) containing the cell annotations. For the first input file, we accept both rdata and plain text file (txt) formats. For rdata inputs, it should contain a seurat object (Satija, et al., 2015). For txt inputs, the single-cell gene expression matrix is in a cell-by-gene format, where each row represents a cell and each column represents a gene, with numerical values indicating the gene expression raw counts. For the second input file, it corresponds to the cell annotations of the gene expression matrix, requiring the inclusion of cell type information for each cell. We also offer users options to specify the minimum number of genes, minimum number of cells, maximum iterations, number of ligand-receptor pairs shown in display, weights for integration, and $p$-value threshold for communication network inference. The corresponding parameter names, descriptions, and default values are provided in Supplementary Table 3.

Upon submission of an inference task, users will receive a task ID immediately. This unique identifier facilitates retrieval of inference results. Additionally, we provide an exemplary task ID for illustration. For tasks involving larger input files, users have the option to provide their email address. Upon completion of the task, an email containing a download link for the inferred results will be automatically dispatched to the user.

**Output**

After setting parameters and submitting the input file, users can obtain the current status and information according to the taskID. The inference results include a table containing information on significant ligand-receptor pairs, an integrated cell communication circular plot, statistic measures of the cell type information, the most frequently occurring ligands and receptors, and heatmaps of ligand-receptor pair interactions. In the visualization section, an integrated cell communication circular plot is displayed, along with cell type information in the dataset, the top most frequently occurring ligands and receptors, and bar plots and heatmaps of ligand-receptor pair interactions. We have also meticulously crafted a tabular format for ease of reference. The rows within this table correspond to the number of significantly expressed ligand-receptor pairs between each pair of cell types, as determined by the specified *p*-value threshold. The table is structured with seven columns, which delineate the inference method, task ID, ligand name, receptor name, receptor-ligand pair name, the calculated communication score, and the *p*-value. Notably, both the ligand and receptor entries are hyperlinked to their respective GeneCards entries. This table is designed to be downloadable, empowering users to integrate the data into their practical applications and research endeavors.

**Intuitively browsing**

The *Browse* page of the collectNET is meticulously organized according to organ systems, offering users a comprehensive and visually engaging experience. Upon the *Home* page, icons representing various organs are readily accessible, and users can delve into detailed cellular landscapes with a single click. For each organ, collectNET presents a nuanced depiction of cellular composition through circular charts that enumerate the cell counts for different cell types. Additionally, collectNET provides a detailed account of the most frequently occurring ligands, receptors, and ligand-receptor pairs, as identified by our cell communication inference methodology, along with their respective frequencies, in a tabular format that underscores the significance of these interactions.

Each organ may encompass one or multiple batches, with each batch representing a distinct dataset. collectNET further enriches the exploration of the user by incorporating UMAP plots, CellChat (Jin, et al., 2021), CellTalker (Cillo, et al., 2020), and CellPhoneDB (Efremova, et al., 2020) inference results, as well as the outcomes of our proprietary cell-cell communication inference method. These are presented alongside a comprehensive table of all significant ligand-receptor pairs within the dataset, offering a multifaceted view of cellular interactions.

The communication networks are elegantly visualized through circular diagrams that encapsulate the quantity of cells within each cell type, the number of significant pathways between different cell types, and the communication score values. This visual representation not only enhances the user's understanding of the cellular ecosystem but also facilitates the identification of key communication hubs and potential targets for therapeutic intervention.

**Advanced searching**

collectNET offers advanced search capabilities that facilitate comprehensive exploration of biological interactions. Users have the flexibility to index and query the database using various parameters such as methods, organs, cell types, receptors, ligands, or ligand-receptor interactions. The system supports multi-criteria indexing, allowing for nuanced searches that can be further refined based on the specific research interests of the user. The search results can be sorted in ascending or descending order according to the *p*-value of the integration method or communication score, providing users with a ranked list of interactions that are statistically significant or biologically relevant. Additionally, the platform utilizes cookies to remember and retain the preferred settings of the user. The search outcomes are also downloadable, enabling users to analyze and integrate the data into their own research workflows seamlessly.

### Text S4. Case applications on the usage of collectNET

**collectNET demonstrates a diverse range of capabilities in new data mining.**

A researcher, intrigued by the characteristics of a particular gene in the context of drug targeting, seeks to understand the signaling pathways in which this gene plays a role, the human organs in which it is most likely involved in cell-cell communication, and the nature of its function. collectNET can substantially aid in this endeavor, as the Fig. S1 shows. Through the *Search* page of collectNET, the researcher can input the name of the ligand or receptor and select to sort the search results by *p*-value or communication score in descending order. The comprehensive search outcomes facilitate a multitude of analytical possibilities for the researcher. Beyond identifying the key cell-cell communication pathways that are associated with a gene of interest, the researcher can delve into the intricacies of these interactions across diverse cell types. The communications are meticulously ranked, providing a clear hierarchy based on the strength of interaction or the level of confidence, which is instrumental in prioritizing further research efforts. Moreover, the search results offer a wealth of ancillary information that enhances the understanding of each ligand-receptor pair. This includes detailed annotations sourced from reputable external databases, which may encompass biological pathways, functional associations, and disease relevance. Such supplementary data points enrich the narrative of molecular interactions and contribute to a more nuanced interpretation of the gene's role in cellular communication. Recognizing the importance of data accessibility, the platform also enables the researcher to download the collected information, allowing for seamless integration into their research workflow. This feature ensures that the data can be utilized within the researcher's preferred analytical tools or stored for future reference. In summary, the search outcomes not only provide a detailed view of the gene's involvement in cell-cell communications but also serve as a versatile resource for exploring the broader biological context, facilitating a deeper and more comprehensive exploration of cellular interactions and functions. Should the researcher have a particular interest in the communication status within certain human organs, the *Browse* page of collectNET allows for a more detailed examination of the information of the organ and statistical data within the atlas.

**collectNET corroborates the efficacy of the inference methodology through statistical methods.**

Owing to the integration of three distinct inference methods in collectNET, we can empirically assess the significance of this integration through statistical methods. We calculated Pearson correlation coefficients (PCC) (Cohen, et al., 2009) on the communication networks inferred by the four methods across ten independent single-cell datasets from HCL (Han, et al., 2020). The PCC is a statistic used to measure the strength and direction of the linear relationship between two variables. In this context, it aims to evaluate the degree of correlation between the two communication networks inferred by different methods.

As shown in Fig. S2, the x-axis represents the three inference methods being compared to the method mentioned in the title, while the y-axis denotes the PCC, where a larger value indicates a higher correlation. Fig. S2A presents a boxplot of the PCC between CellChat and the other methods, with the median value of collectNET 0.77 and markedly higher than the other methods. This indicates a strong correlation between the results of collectNET and those of CellChat, surpassing the correlations observed between CellChat and the other two methods. Similarly, Fig. S2B and S2C demonstrate that, the results of collectNET are highly correlated to CellPhoneDB and CellTalker, with the median value 0.87 and 0.99 respectively. One-sided Wilcoxon signed-rank test (Woolson, 2007) were conducted on the PCCs of collectNET and the second-ranked method, and the overall PCC of collectNET is significantly higher than each of the second-ranked method, with *p*-values 0.006, less than 1e-16 and less than 1e-16 respectively. This underscores that despite the obvious differences between the networks inferred by different methods, collectNET effectively integrates the methods based on different models and databases of ligand-receptor pairs, constructing a more accurate communication network by combining diverse biological information.

**collectNET reveals the topological characteristics of communication networks.**

The communication network inferred by collectNET can be regarded as a two-dimensional directed graph with weighted edges, where the weights are represented by communication scores. The topological information within biological networks, including node information and subgraph information, reveals the existence of pivotal cell types and potential distinctive patterns within cellular signaling.

As an algorithmic measure of the relative importance of a node within a network, the PageRank score is used to explore the role assumed by each node. The PageRank algorithm can be expressed by the following formula for iterative computation.

$$P_i^{(t+1)} = \alpha \sum_{j \in N(i)} \frac{P_j^{(t)}/L_j}{N_j} + (1-\alpha)\frac{1}{N}$$

where $P_i^{(t)}$ is the PageRank value of cell type $i$ at iteration $t$, $\alpha$ is the damping factor set to 0.85, $L_j$ is the total number of outgoing links from cell type $j$, and $N$ is the total number of cell types in the network. At the start of the iterative algorithm, the PageRank value of each node is set to $1/N$. The iteration stops after multiple iterations when the change in PageRank values between two consecutive iterations is less than a certain threshold, which is set at 1% of the initial value. Within communication networks provied by collectNET in the tutorial data, the PageRank score quantifies the significance of interactions within cell type pairs throughout the biological process. As shown in Fig. S3, the ventricle cardiomyocyte exhibits a PageRank score approaching 0.2. This corresponds to the ventricle cardiomyocytes as a specialized class of myocytes that are uniquely found within the ventricles of the heart and play a significant role in cardiac regeneration, development, cell cycle regulation, and electrophysiological properties. In contrast, dendritic cells, endothelial cells, and neutrophils all exhibit significantly lower scores, suggesting a more singular signaling pathway presence within these cell types.

The largest clique in a graph is a subset of vertices where each pair is adjacent and no more vertices can be added (Abello, Pardalos and Resende, 1998), and in biological networks it is often used to identify common structural patterns or motifs that occur frequently, and thus provide valuable insights into the mechanisms of cellular processes, disease pathways, and evolutionary relationships. By utilizing iterative algorithms with a backtracking method, we have pinpointed the largest clique within a communication network. The network provided by collectNET has revealed that ventricle cardiomyocytes, macrophages, fibroblasts, smooth muscle cells, M1 and M2 macrophages form this prominent clique, indicating extensive shared communication patterns and active interactions.

Moreover, the graph clustering analysis of the network, which assesses edge connectivity through betweenness, has identified two distinct clusters: one with endothelial cells and neutrophils, and another with dendritic cells. Since endothelial and neutrophil interactions are crucial in inflammation and immune responses, including the release of inflammatory mediators and neutrophil migration, highlighting their functional interplay in cardiovascular and immunological dynamics (Phillipson and Kubes, 2011; Tonnesen, 1989), this segregation is biologically significant.

#### Text S5. System design and implementation

collectNET website is maintained on a Linux-based Apache web server v2.4.58 (http://www.apache.org), while optimizing the web interface with the Bootstrap v3.3.7 (https://getbootstrap.com/docs/3.3/) framework. To implement advanced tables and responsive charts, we utilize a range of jQuery Plugins and JavaScript Libraries, including DataTables v1.10.19 (https://datatables.net). The current version has ensured compatibility with mainstream web browsers such as Google Chrome, Firefox, Opera, Microsoft Edge, Apple Safari, etc.

### Supplementary Figures

#### Fig. S1. collectNET demonstrates a diverse range of capabilities in new data mining.

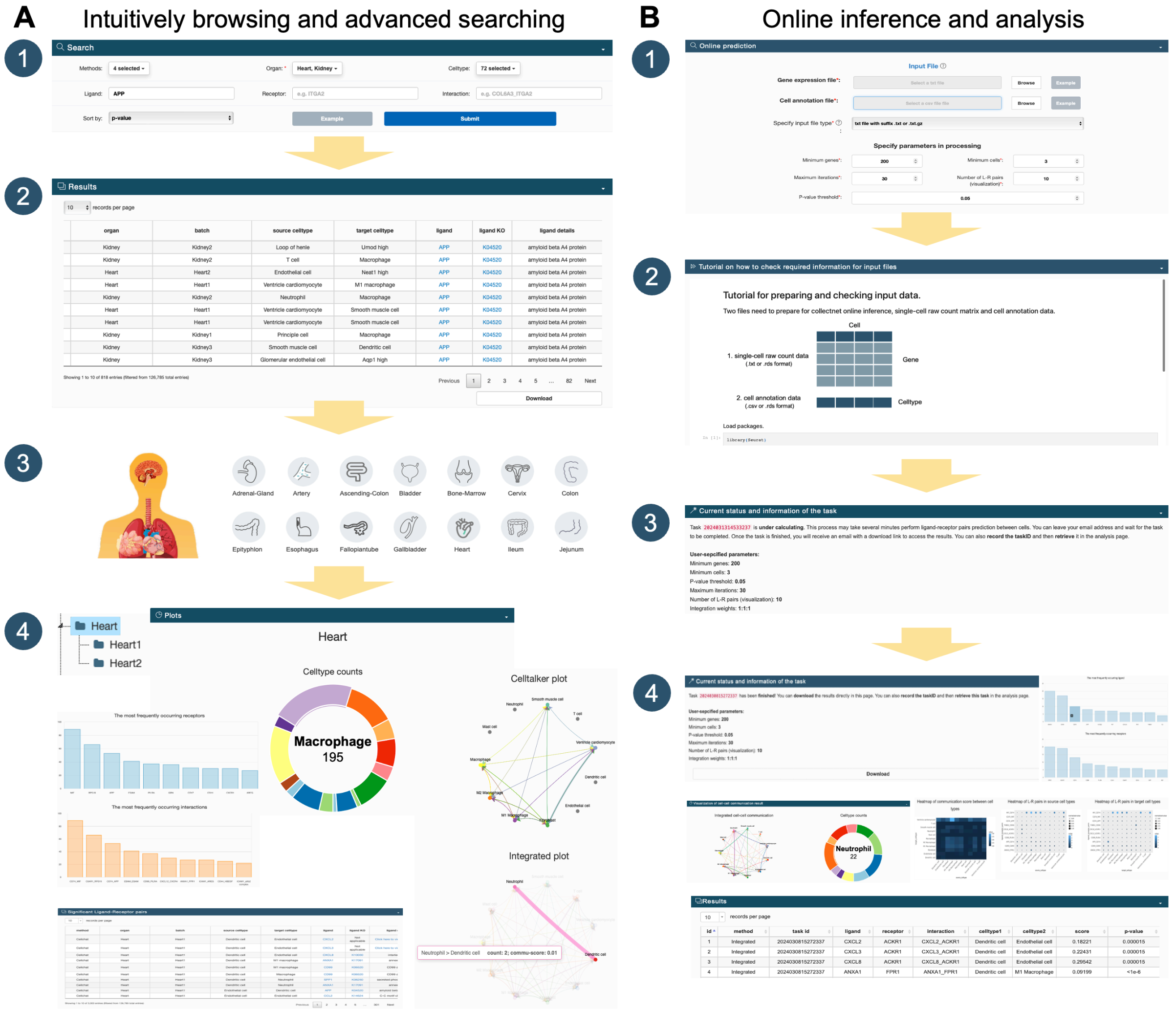

**Fig. S1. collectNET demonstrates a diverse range of capabilities in new data mining.** collectNET helps to understand the signaling pathways in which this gene plays a role, the human organs in which it is most likely involved in cell-cell communication, and the nature of its function.

**Fig. S2. collectNET corroborates the efficacy of the inference methodology through statistical methods.**

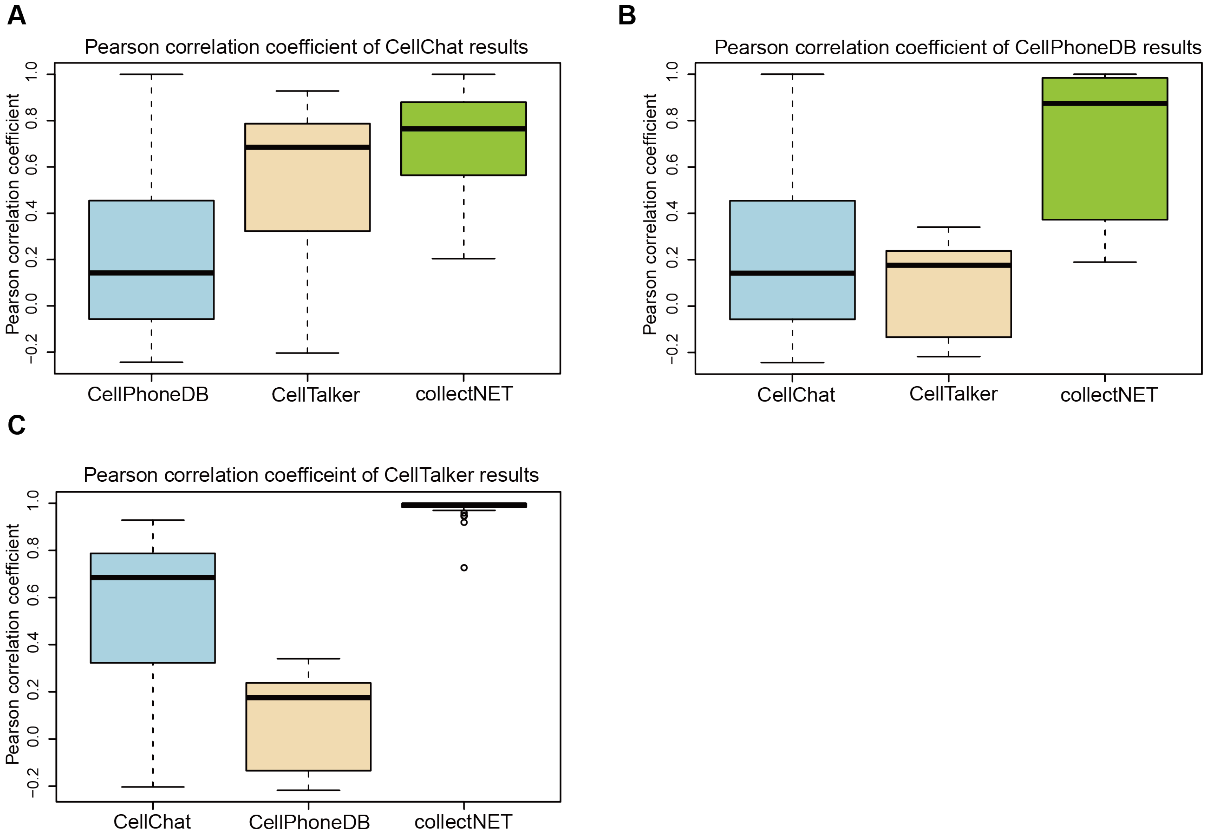

**Fig. S2. collectNET corroborates the efficacy of the inference methodology through statistical methods.** (A) Boxplot of the Pearson correlation coefficient between the CellChat method and the other methods. (B) Boxplot of the Pearson correlation coefficient between the CellPhoneDB method and the other methods. (C) Boxplot of the Pearson correlation coefficient between the CellTalker method and the other methods.

**Fig. S3. collectNET reveals the topological characteristics of communication networks.**

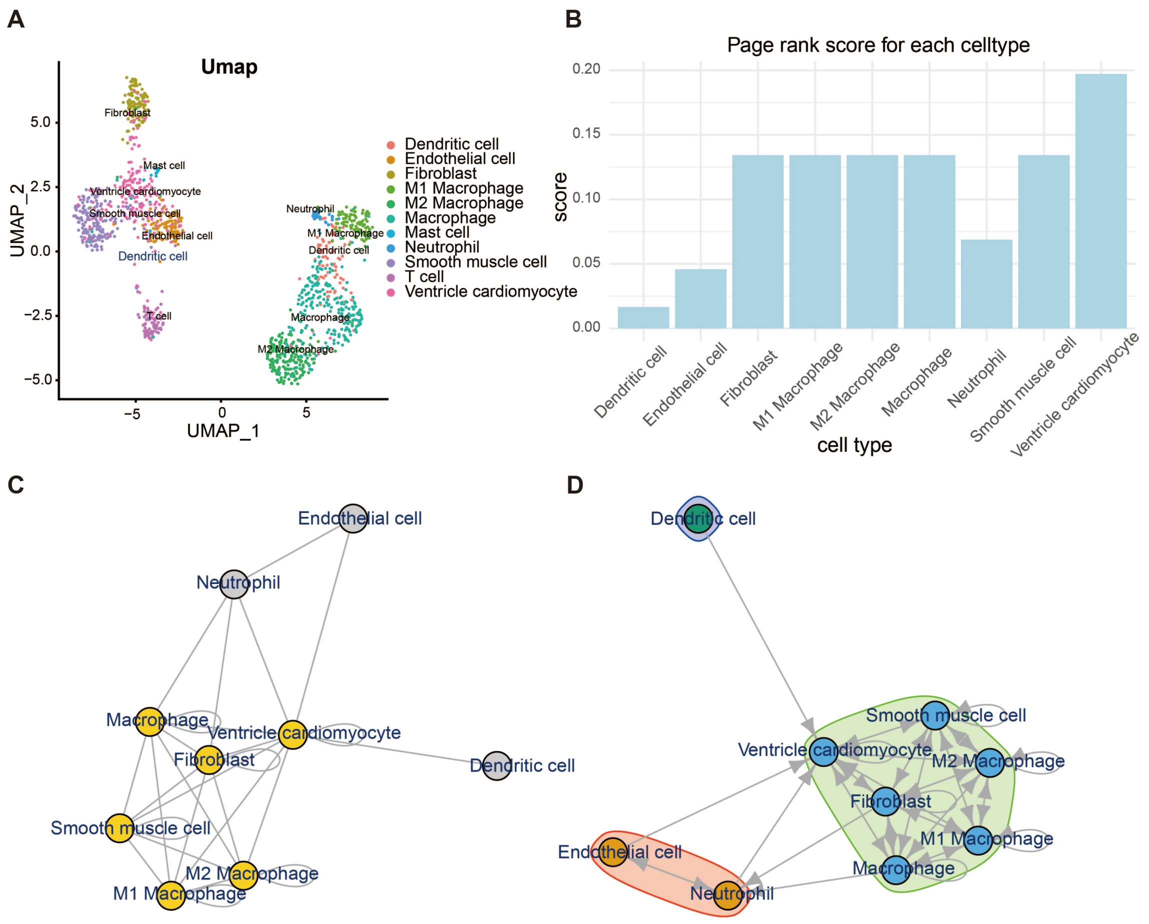

**Fig. S3. collectNET reveals the topological characteristics of communication networks.** (A) Umap plot on tutorial data. (B) PageRank score quantifies the significance of each cell type's interactions with others throughout the biological process. (C) The left side shows the largest clique found by using graph search algorithms, while the right side shows the graph clustering based on strong connectivity. Both provide insights into the functional relationships and potential collaborative roles of these cell types.

#### Fig. S4. Computational efficiency of collectNET.

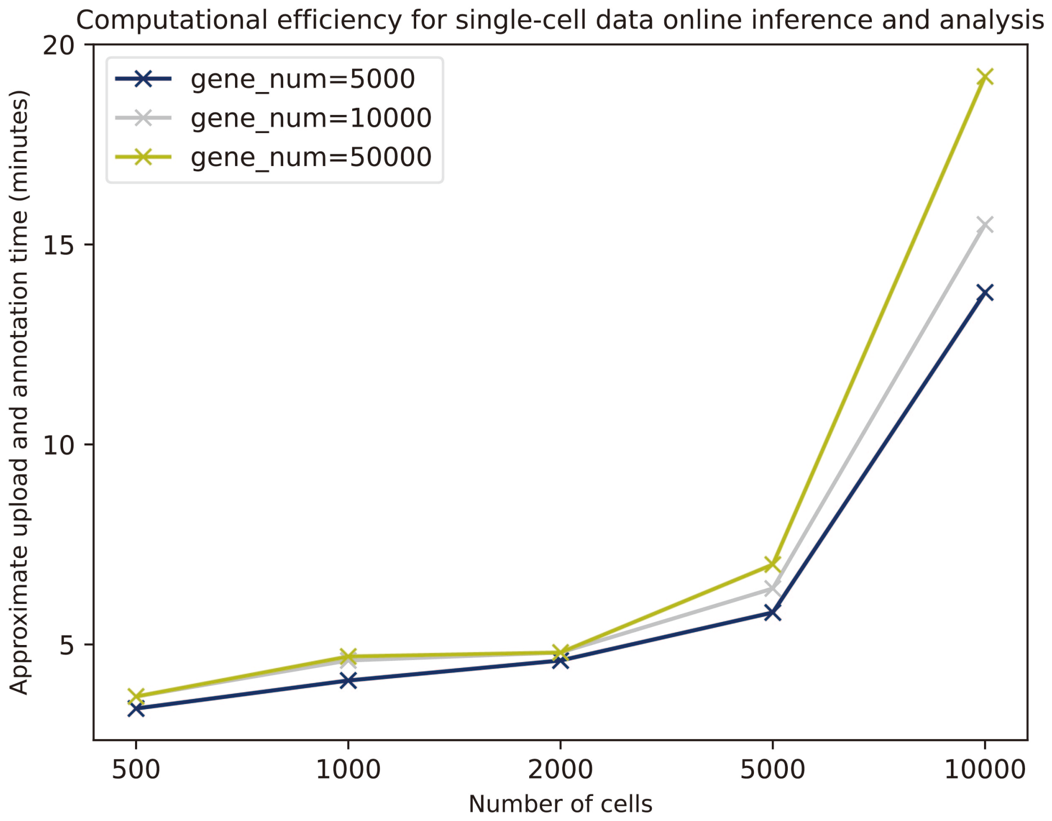

**Fig. S4. Computational efficiency of collectNET.** Computational efficiency for single-cell data online inference and analysis, where the x-axis denotes the number of cells ranging from 500 to 10,000, and the y-axis denotes approximate upload and annotation time (minutes).

#### Supplementary Tables

**Table S1.** Comparison of collectNET with other published cell-cell communication databases or websites

| **Categories** | **Functionalities and applications** | **collectNET** | **CellCommuNet (Ma, et al., 2024)** | **CITEdb (Shan, et al., 2022)** | **TALKIEN (Moratalla-Navarro, Moreno and Sanz-Pamplona, 2023)** |
|---|---|---|---|---|---|
| Prior data | Integration of various L-R pair databases | ✓ | ✓ | | ✓ |
| Methods for inference | CellChat | ✓ | ✓ | | |
| | CellPhoneDB | ✓ | | | |
| | CellTalker | ✓ | | | |
| Database | Diverse forms of statistical graphs | ✓ | ✓ | ✓ | ✓ |
| | Multiple externel hyperlinks | ✓ | ✓ | ✓ | ✓ |
| | Multi-tiered Search and Download funtionality | ✓ | ✓ | ✓ | |
| Web service | Support for user-provided single-cell data | ✓ | ✓ | | |
| | Innovative inference methology | ✓ | | | |
| | Customizable parameters for network construction | ✓ | | | |
| | Downstream analysis of the network | ✓ | | | ✓ |

**Table S2.** Comparison of collectNET and other reference ligand-receptor pair databases

| L-R Databases | L-R pairs | Contain Copolymers |
|---|---|---|
| collectNET | 3954 | Yes |
| CellChatDB | 1939 | Yes |
| CellPhoneDB | 1396 | Yes |
| CellTalkDB | 2557 | No |

**Table S3.** The user-defined parameters for collectNET.

| **Parameters** | | **Description** |
|---|---|---|
| Gene expression file | | The input single-cell RNA sequencing matrix file, both txt and rds formats supported. |
| Cell annotation file | | The input single-cell data cell labels, csv format supported. |
| Specify input file type | | The format of the input file. |
| Minimum genes | | The minimum number of genes required per cell (for cell filtering). |
| Minimum cells | | The minimum number of cells in which a gene must be expressed (for gene filtering). |
| Maximum iterations | | The number of iterations for inference in CellPhoneDB and CellTalker methods. |
| Number of L-R pairs (visualization) | | The most frequently occurring ligand-receptor pairs and their quantities displayed in the result visualization. |
| *p*-value threshold | | The threshold for L-R pairs considered significant in the inference results. |
| Weight for integration | CellChat | The weight of communication values in the CellChat method within the integrated approach. |
| | CellPhoneDB | The weight of communication values in the CellPhoneDB method within the integrated approach. |
| | CellTalker | The weight of communication values in the CellTalker method within the integrated approach. |
